# Transcriptome sequencing suggests that pre-mRNA splicing counteracts widespread intronic cleavage and polyadenylation

**DOI:** 10.1101/2022.05.27.493724

**Authors:** Mariia Vlasenok, Sergey Margasyuk, Dmitri D. Pervouchine

**Affiliations:** Skolkovo Institute of Science and Technology, Moscow 143026, Russia

**Keywords:** alternative splicing, intronic polyadenylation, polyadenylated introns, APA, PAS

## Abstract

Alternative splicing (AS) and alternative polyadenylation (APA) are two crucial steps in the post-transcriptional regulation of eukaryotic gene expression. Protocols capturing and sequencing RNA 3’-ends have uncovered widespread intronic polyadenylation (IPA) in normal and disease conditions, where it is currently attributed to stochastic variations in the pre-mRNA processing. Here, we took advantage of the massive amount of RNA-seq data generated by the Genotype Tissue Expression project (GTEx) to simultaneously identify and match tissue-specific expression of intronic polyadenylation sites with tissue-specific splicing. A combination of computational methods including the analysis of short reads with non-templated adenines revealed that APA events are more abundant in introns than in exons. While the rate of IPA in composite terminal exons and skipped terminal exons expectedly correlates with splicing, we observed a considerable fraction of IPA events that lack AS support and attributed them to spliced polyadenylated introns (SPI). We hypothesize that SPIs represent transient byproducts of a dynamic coupling between APA and AS, in which the spliceosome removes an intron after cleavage and polyadenylation have already occurred in it. These findings indicate that cotranscriptional pre-mRNA splicing could serve as a rescue mechanism to suppress premature transcription termination at intronic polyadenylation sites.

## Introduction

The majority of transcripts that are generated by the eukaryotic RNA Polymerase II undergo endonucleolytic cleavage and polyadenylation (CPA) at specific sites called the polyadenylation sites (PASs) (*1*). More than half of human genes have multiple PASs resulting in alternative polyadenylation (APA) (*2, 3*). APA modulates gene expression by influencing mRNA stability, translation, nuclear export, subcellular localization, and interactions with microRNAs and RNA binding proteins (RBPs) (*4, 5*). APA is widely implicated in human disease, including hematological, immunological, neurological disorders, and cancer (*6, 7*).

APA can generate transcripts not only with different 3’-untranslated regions (3’-UTR) but also transcripts encoding proteins with different C-termini (*8*). Recent studies have shown that more than 20% of human genes contain at least one intronic PAS located upstream of the 3’-most exon (*9*). Intronic polyadenylation (IPA) can lead to important functional changes due to alterations in the protein primary sequence (*10*). For instance, IPA in DICER generates a truncated protein with impaired miRNA cleavage ability that results in decreased endogenous miRNA expression (*11, 12*). Remarkably, the truncated oncosuppressor proteins that are generated by IPA often lack tumor-suppressive functions and contribute significantly to the tumor onset and progression (*11*).

The interplay between splicing and polyadenylation has long been recognized as being related to cotranscriptional pre-mRNA processing (*13*). Many splicing factors have dual roles serving both splicing and polyadenylation, including *U2AF* (*14*), *PTBP1* (*15*), members of Hu protein family (*16*), and others (*8*). The observation that IPA is associated with weaker 5’-splice sites and longer introns, and experiments on mutagenesis of CPA and splicing signals in plants together suggest that splicing and polyadenylation operate in a dynamic competition with each other (*9, 17*). Furthermore, nascent RNA polymerase II transcripts that are susceptible to CPA at cryptic PASs are protected from CPA by U1 snRNP in a process called telescripting, most remarkably in genes with longer introns (*18*). These results raise a number of challenging questions about the actual abundance and function of cryptic intronic PASs, mechanisms of their inactivation, and relation to AS.

A number of experimental protocols have been developed to identify the genomic positions of PASs (*19*). Many of them use oligo(dT) (3’RNA-seq, PAS-seq, polyA-seq) or similar primers (3’READS) to specifically capture transcript ends (*20–23*). A combination of these protocols yielded a consolidated set of more than 500,000 human PASs (*24–26*), however many more PASs may be active in tissue- and disease-specific conditions. A number of computational methods also attempt to identify PASs from the standard polyA^+^ RNA-seq data as genomic loci that exhibit an abrupt decrease in read coverage (*27–32*). However, since the density of RNA-seq reads is highly non-uniform along the gene length, many of these methods are limited to PASs that are located in the last exon or 3’-UTR, thus implicitly focusing on quantifying relative usage of PASs in the gene 3’-end rather than on identifying novel intronic PASs.

On the other hand, RNA-seq data contain an admixture of reads that cover the junction between the terminal exon and the beginning of the polyA tail. They align to the reference genome only partially due to a stretch of non-templated adenine residues. Although the fraction of such reads is quite small and normally does not exceed 0.1%, they can potentially be used for *de novo* identification of PASs. Previous studies such as ContextMap2 (*31*) and KLEAT (*30*) demonstrated that the analysis of RNA-seq reads containing a part of the polyA tail can offer a powerful alternative to coverage-based methods when analyzing a sufficiently large panel of RNA-seq experiments.

In this work, we took advantage of the massive amount of RNA-seq data generated by the Genotype Tissue Expression Project (GTEx), the largest to-date compendium of human transcriptomes, to simultaneously assess alternative splicing and intronic polyadenylation and match their tissue-specific patterns (*33*). Unlike previous studies, which extensively characterized the tissue-specific polyadenylation using coverage-based methods (*28, 34–36*), here we focused specifically on intronic PAS by combining the information on polyA reads to identify PAS, split reads to measure the AS rate, and the read coverage to assess the CPA rate. We identified a core set of 318,898 PAS clusters that are stably expressed in GTEx tissues, which is consistent with other published sets, and characterized their attribution to the UTRs, exonic, and intronic regions. After normalizing the number of polyA reads to the background read coverage in exons and introns, we observed that intronic PAS are used more frequently than PASs in regions that are not spliced, i.e., exons. Moreover, in inspecting the concordance between IPA and AS, we unexpectedly found a considerable fraction of unannotated intronic PAS that are inconsistent with previously proposed CPA models. We attributed them to Spliced Polyadenylated Introns (SPI), a term we introduce here to describe transient byproducts of the dynamic coupling between CPA and AS, and conjecture that they are generated when the spliceosome removes an intron after CPA have already occurred in it.

## Results

### The identification of PAS

The majority of short reads in the output of polyA^+^ RNA-seq protocols align perfectly to the genome, but a small fraction map partially due to stretches of non-templated adenines generated by CPA. Since RNA-seq reads with incomplete alignment to the genomic reference tend to map to multiple locations, we took a conservative approach by analyzing only uniquely mapped reads from 9,021 GTEx RNA-seq experiments (*33*) with additional restrictions on sequencing quality (see Methods). We extracted polyA reads, defined as short reads containing a soft clipped region of at least six nucleotides that consists of 80% or more adenines, excluding reads aligning to adenine-rich genomic tracks and omitting samples with exceptionally large numbers of polyA reads (Figure S1). Out of ∼ 356 billion uniquely mapped reads, ∼ 591 million (0.17%) polyA reads were obtained. At that, the average adenine content in soft clipped regions of polyA reads was 98% despite the original 80% threshold, confirming that the selected short reads indeed contain polyA tails.

The alignment of a polyA read is characterized by the genomic position of the first non-templated nucleotide, which presumably corresponds to a PAS, and the length of the soft clip region, referred to as overhang (Figure 1A). Consequently, each PAS is characterized by the number of supporting polyA reads, referred to as polyA read support, and the distribution of their overhangs. Our confidence in PAS correlates not only with polyA read support, but also with the diversity of the overhang distribution, which is measured by Shannon entropy *H*. Out of 9.6 million candidate PASs, 2.1 million (22%) had *H* ≥ 1 and 565,387 (6%) had *H* ≥ 2 (Figure S2). In further analysis, we chose to use the threshold *H* ≥ 2 in order to obtain a list of PASs that matches by the order of magnitude the consolidated atlas of polyadenylation sites from 3’-end sequencing (*24*) and captures sufficiently many annotated gene ends (Supplementary File 1). Out of 565,387 PASs with *H* ≥ 2, 331,563 contained a sequence motif similar to the canonical consensus CPA signal (NAUAAA, ANUAAA, or AAUANA) in the 40-nt upstream region (*37, 38*). The latter PASs will be referred to as PASs with a signal.

**Figure 1:**
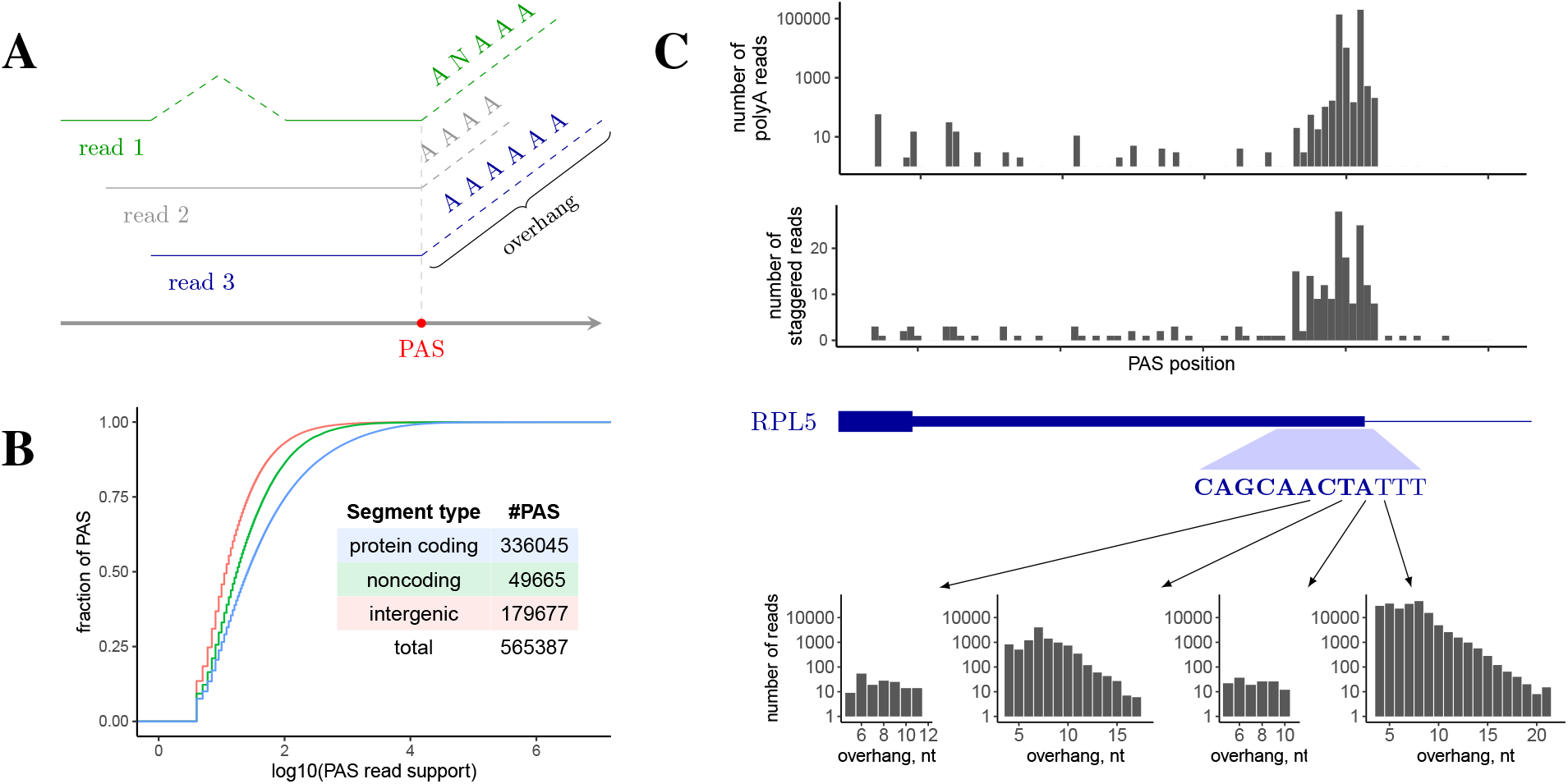
The identification of PAS. **(A)** The alignments of short reads with non-templated adenine-rich ends (polyA reads). The genomic position of the first non-templated nucleotide corresponds to a PAS. The length of the soft clip region is referred to as overhang. **(B)** The polyA read support of PAS in protein-coding genes, non-coding genes, and intergenic regions. The number of PASs in each group is indicated in the inset. **(C)** The 3’-end of the *RPL5* gene is highly covered by polyA reads. Top: the positional distribution of the number of polyA reads and the number of staggered polyA reads (i.e., the number of different overhangs). Bottom: the distribution of overhangs at the indicated positions.

To characterize the occurrence of PASs in different genomic regions, we subdivided the human genome into a disjoint union of intervals corresponding to protein-coding genes, non-coding genes, and intergenic regions. In total, 336,045, 49,665, and 179,677 PASs were detected in these respective regions; of these 69%, 61%, and 39% were PASs with a signal, respectively. The level of polyA read support in different genomic regions also varied, e.g. 25.5%, 14%, and 7% PASs were supported by 100 or more polyA reads in protein-coding, non-coding, and intergenic regions, respectively (Figure 1B). As expected, protein-coding regions had the largest density of PASs per megabase. However, a large absolute number of PASs in intergenic regions, including PASs without canonical consensus CPA signals, indicates that a substantial number of RNA Pol II transcripts are transcribed from them, in accordance with current hypotheses on pervasive transcription (*39–41*).

An example of a gene that is highly covered by polyA reads is *RPL5* (Figure 1C). We identified several PASs in the vicinity of its annotated transcript end (TE), some of which were supported by as many as 100,000 polyA reads with more than 20 different overhangs. Unexpectedly, instead of a single peak, we observed a relatively dispersed cluster of PASs spanning twelve nucleotides. Manual inspection confirmed that the RNA-seq read alignments ending in all these positions indeed were followed by non-templated polyA tracks, thus indicating that the observed pattern is due to biological stochasticity and not due to mapping artifacts. Remarkably, the number of polyA reads decayed with increasing the length of the overhang (Figure 1C, bottom). This decrease could result from the mapping bias, in which a lower fraction of reads with larger soft clip regions can be mapped uniquely, or be a consequence of degradation of the substrates possessing multiple terminal adenines by exonucleases (*42*).

### PAS clusters

Large variability of PASs positions in *RPL5* motivated us to explore the distribution of distances from each PAS to its closest annotated TE in protein-coding genes (Figure 2A). Among PASs that were located within 100 nts from an annotated TE, 71% fell within 10 nts, and 78% of PASs with a signal did so. We therefore chose to cluster PASs that were located within 10 nts of each other (Figure 2B). This yielded 318,898 PAS clusters (PASCs), of which 90% had length below or equal to 10 nts, 72% consisted of a unique PAS, and 99% consisted of less than ten individual PASs (Supplementary File 2). In what follows, a PASC will be referred to as PASC with a signal if it contains at least one individual PAS with a signal; the polyA read support of a PASC is defined as the total number of supporting polyA reads of its constituent individual PASs.

**Figure 2:**
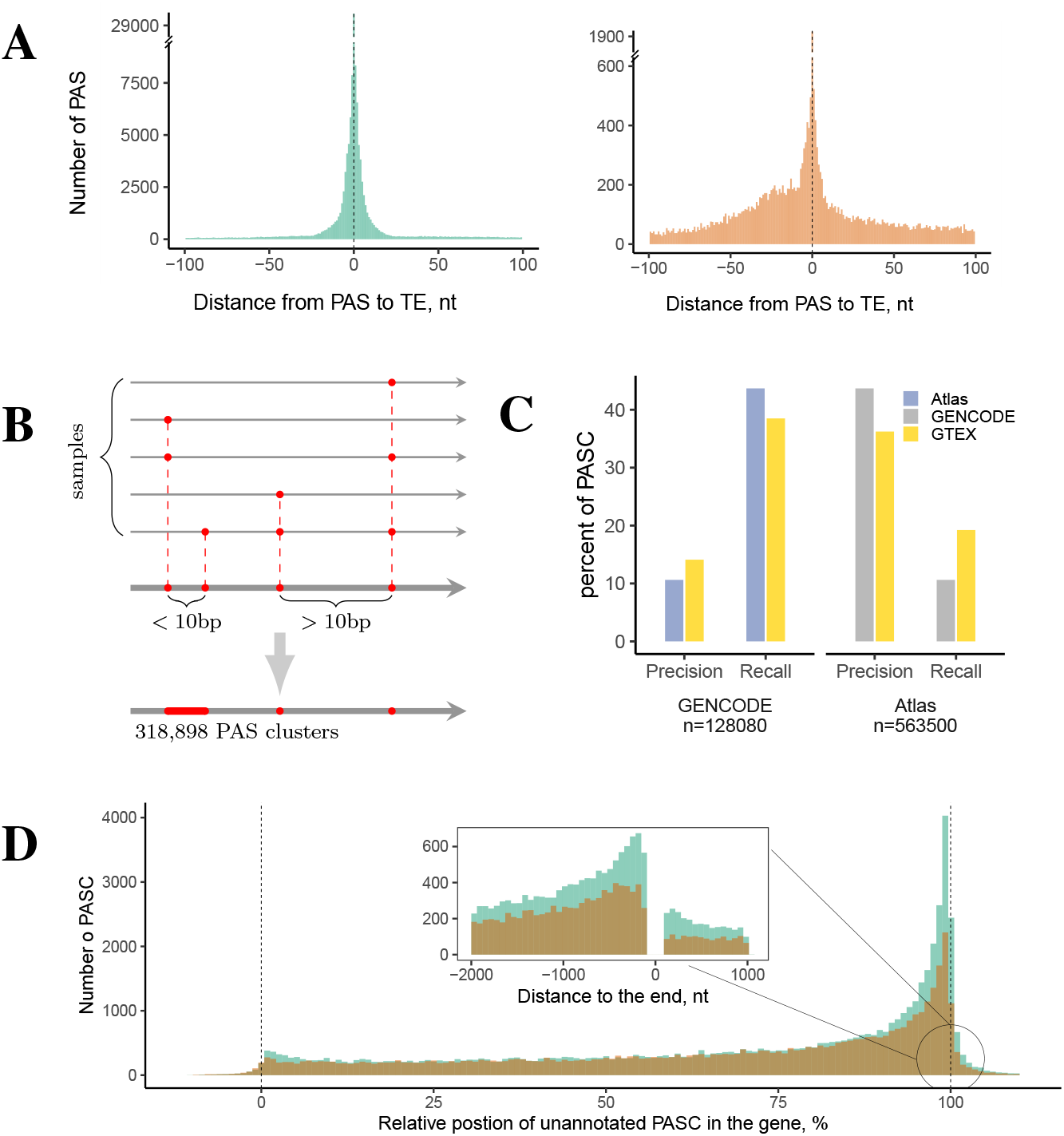
PAS clusters in protein-coding genes. **(A)** The distribution of distances from each PAS to its closest annotated transcript end (TE) for PAS with (*n* = 122, 448) and without a signal (*n* = 22, 361). **(B)** PAS located *<*10 bp from each other are merged into PAS clusters (PASCs). **(C)** Pairwise comparison of PASs inferred from GTEx, PolyASite 2.0 (*24*) (Atlas), and GENCODE. Left: the proportion of PASC from GENCODE that are supported by Atlas or GTEx (precision) and the proportion of PASC from Atlas or GTEx that are supported by GENCODE (recall). Right: the proportion of PASC from Atlas that are supported by GENCODE or GTEx (precision) and the proportion of PASC from GENCODE or GTEx that are supported by Atlas (recall). **(D)** The relative positions of unannotated PASCs (i.e., ones not within 100 bp of any annotated TE) along the gene length. 0% and 100% correspond to the 5’-end and 3’-end of the gene, respectively. The inset shows distribution of absolute positions of unannotated PASCs around the gene end.

We next asked how PASCs identified from GTEx RNA-seq data correspond to the consolidated polyadenylation atlas derived from 3’-end sequencing (PolyASite 2.0 (*24*), in what follows referred to as Atlas) and TEs annotated by the GENCODE consortium (*43*). To assess this, we surrounded TEs from GENCODE by 100-nt windows and analyzed pairwise intersections of the three respective sets (Figure 2C). The precision of GTEx with respect to GENCODE, i.e., the proportion of PASCs from GTEx that were located within 100 nts of an annotated TE, was higher than that of PolyASite 2.0, while the recall, i.e., the proportion of annotated TEs that are supported by at least one PASC from GTEx within 100 nts, was lower. Conversely, the precision of GTEx with respect to PolyASite 2.0 was lower compared to that of GENCODE, while the recall was higher. This comparison indicates that GTEx RNA-seq data yields a slightly more conservative set of PASCs than PolyASite 2.0. The benefit of using GTEx PASCs is that RNA-seq provides a snapshot of alternative splicing and polyadenylation assessed in the same conditions. Additional analysis of the relationship between precision and recall for GTEx and PolyASite 2.0 weighted by the polyA read support confirmed that the two sets are largely consistent (Figure S3).

Since 85% of newly identified PASCs did not have an annotated TE within 100 nts, we focused on this group of PASCs (referred to as unannotated PASCs) and explored their relative position within the gene length, which is equal to 0% and 100% for the 5’-end and 3’-end of the gene, respectively (Figure 2D). Despite TEs no longer being considered, we observed a considerable increase in PASC density towards the 3’-end for those with and without a signal, and a much weaker, but noticeable increase in the 5’-end. This recapitulates the general tendency of PASCs to occur more frequently towards the 3’-end of the gene, a pattern that is also observed for unannotated PASCs from Atlas (Figure S4).

### PAS clusters in protein-coding regions

We next focused on a subset of 164,497 PASCs that were located in protein-coding genes and explored their attribution to gene parts, namely to the 5’-untranslated region (5’-UTR), the 3’-untranslated region (3’-UTR), and the coding part (CDS). Each CDS region was further sub-divided into intronic, always (constitutive) exonic, and alternative exonic parts (see Methods). Since these regions differ by length, we quantified PASCs not only by absolute number but also by density, i.e., the number of PASCs per nucleotide. Additionally, we quantified the expression of PASCs by taking into account their polyA read support, in which each PASC was weighted by the number of supporting polyA reads (Figure 3).

**Figure 3:**
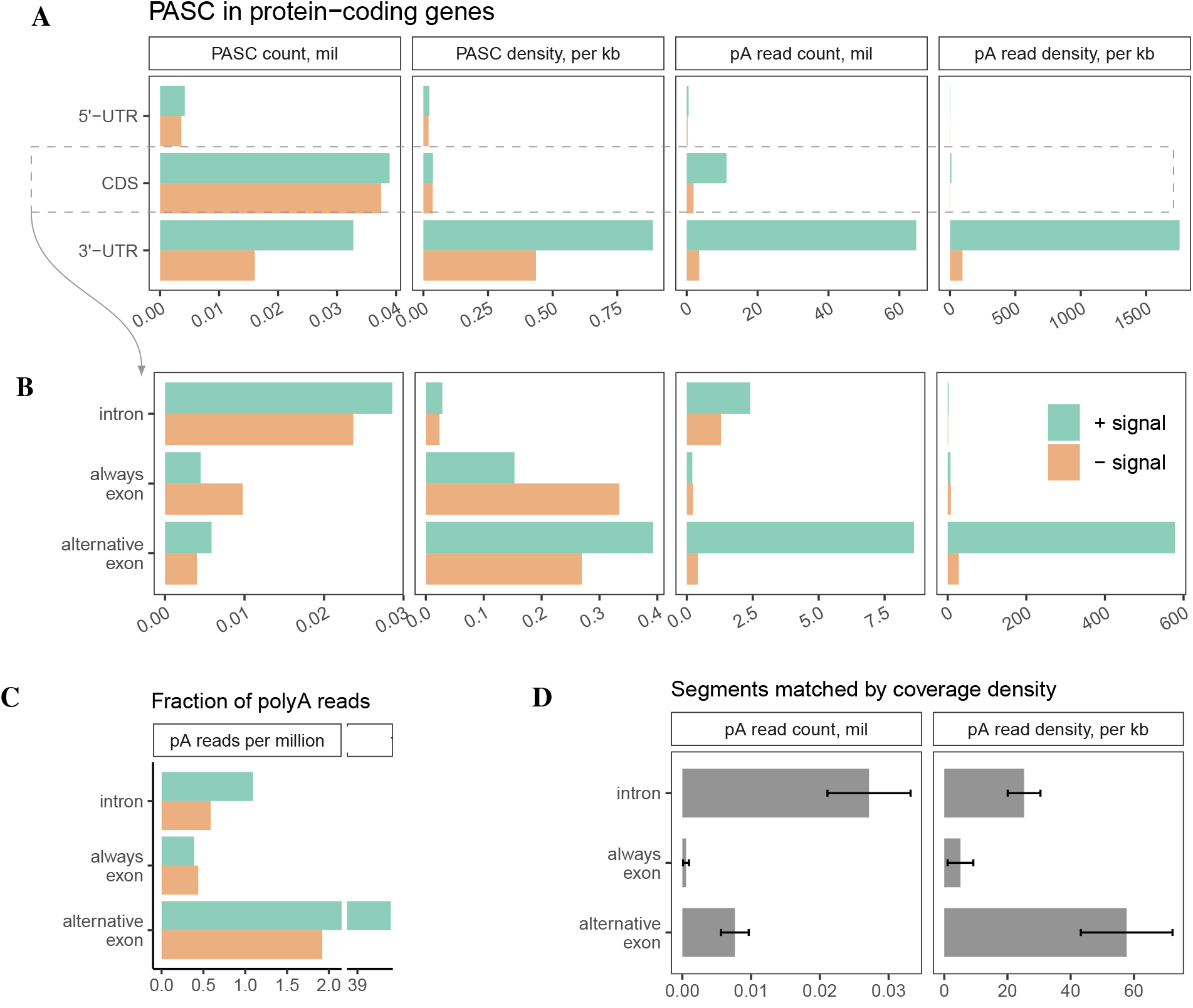
PAS clusters in protein-coding regions. **(A)** The distribution of PASCs in 5’-UTRs, CDS, and 3’-UTRs. Shown are the total number of PASC (PASC count), PASC density per Kb (PASC density), the total number of polyA reads (polyA read count), the total number of polyA reads per kb (polyA read density). **(B)** The distribution of PASCs from CDS in introns, constitutive exons (always exon), and alternative exons. **(C)** The number of polyA reads normalized to the average read coverage in each region (defined as the number of polyA reads per million aligned reads; see Methods for details). **(D)** The number of polyA reads in segments matched by the read coverage density.

As expected, PASCs were quite frequent in CDS by absolute number, but their density was the highest in 3’-UTRs since CDS regions are also longer than UTRs (Figure 3A). The enrichment in 3’-UTRs was more prominent when taking into account the number of supporting polyA reads. Similarly, PASCs were most frequent in introns by absolute number, but their density was the lowest after normalization (Figure 3B). The positional distribution of PASCs had a pronounced peak in the end of exonic regions and in the beginning of intronic regions (Figure S5), and similar peaks were also observed for PolyASite 2.0 (Figure S6). However, despite low density, intronic PASCs were still quite frequent by absolute number, and among them there could be PASCs leading to premature CPA.

Current models assume that introns containing PASs cannot undergo splicing after they are cleaved and polyadenylated (*1*). Here we challenge this assumption by supposing that splicing machinery can operate on introns after CPA have already occurred in them. Such introns would be invisible for RNA-seq after they are removed by the spliceosome and degraded. To estimate the actual CPA rate taking into account this undercoverage bias, we normalized the number of polyA reads to the average read coverage in the respective regions and found that the relative abundance of polyA reads in introns is substantially larger than that in constitutive exons (Figure 3C). Furthermore, we matched introns, constitutive exons, and alternative exons by the read coverage (Figure S8A) and selected a subset of intervals of each type that were covered by approximately the same number of reads (133 *±* 6.7 reads per kb per sample). Then, we computed the number of polyA reads in these matched subsets and, again, found a prominent enrichment of polyA reads in introns as compared to constitutive exons both in terms of the number of polyA reads (Figure 3D, left) and density per nt (Figure 3D, right). Alternative exonic regions expectedly contained a larger number of polyA reads due to the presence of PASs near the endogenous terminal exons. This enrichment remained significant when choosing other read coverage values for matching (Figure S8B,C), in sum indicating that if introns and exons were equally represented in RNA-seq data, the frequency of CPA events in introns would have appeared several times larger than that in constitutive exons.

### Tissue-specific polyadenylation

While PASC positions can be robustly identified by pooling hundreds of millions of polyA reads across the entire GTEx dataset, the rate of their tissue-specific usage cannot be assessed in the same way due to insufficient number of polyA reads in individual samples. Instead, the rate of PASC expression in tissues can be measured by coverage-based methods, as their positions have been already identified. Here, we adapted a simple procedure from (*11*), in which the average read coverage was measured in 150-nt windows, *wi*_1_ and *wi*_2_, before and after each PASC. To quantify PASC expression, we used log_10_(*wi*_1_*/wi*_2_) metric, which captures the magnitude of read coverage drop at a PASC, and a more elaborate method based on DESeq2 (*44*), which additionally accounts for variation between samples (Figure 4A).

**Figure 4:**
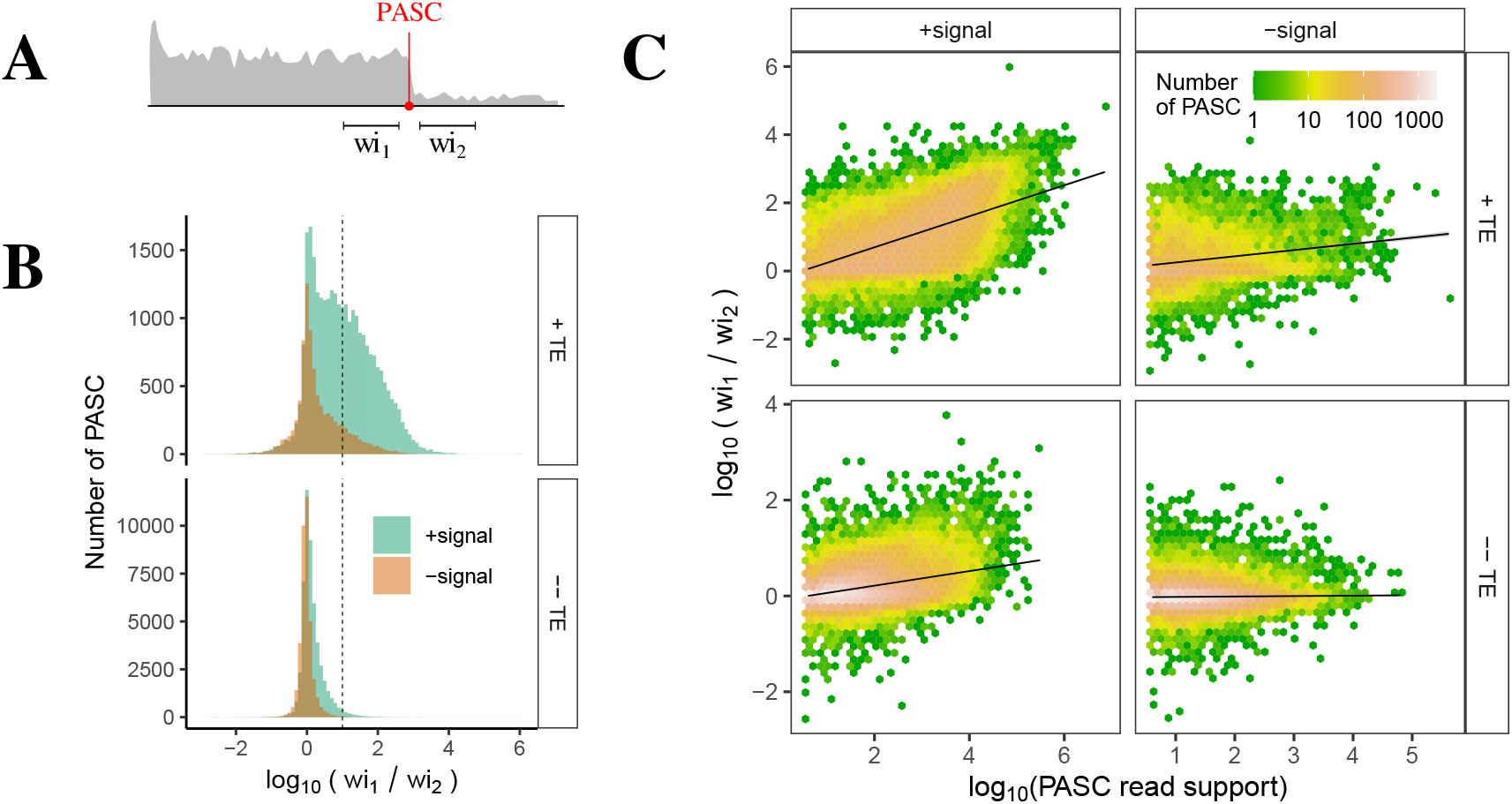
Coverage-based metrics of PASC expression. **(A)** The average read coverage was measured in 150-nt upstream and downstream windows, *wi*_1_ and *wi*_2_, around PASC. **(B)** The distribution of log_10_(*wi*_1_*/wi*_2_) metric for annotated (*n* = 37, 194, top) and unannotated PASCs (*n* = 89, 116, bottom). A PASC is referred to as annotated if it is within 100 bp of an annotated TE. The dashed line represents the cutoff *wi*_1_*/wi*_2_ = 10. **(C)** The log_10_(*wi*_1_*/wi*_2_) metric positively correlates with the number of supporting polyA reads not only for annotated, but also for unannotated PASCs with a signal.

First, we analyzed the set of 164,497 PASCs in protein-coding genes by pooling read coverage profiles across all GTEx samples and excluding PASCs located within 200 nts from splice sites to avoid measuring the read coverage drop at exon-intron boundaries. In the resulting set of 126,310 PASCs (Supplementary File 3), the read density in *wi*_1_ and *wi*_2_ averaged to 8.8 and 3.7 reads per nucleotide per sample, respectively, indicating at least twofold average drop after PASCs. Consistently, the *wi*_1_*/wi*_2_ distribution was skewed towards positive values with a noticeably bigger skewness for PASCs with a signal and PACSs near annotated TEs (Figure 4B). Remarkably, the number of supporting polyA reads was positively correlated with *wi*_1_*/wi*_2_ not only for PASCs near annotated TEs, but also for unannotated PASCs with a signal (Figure 4C). For each PASC, we computed the average read density in *wi*_1_ and *wi*_2_ separately in each tissue. Out of 126,310 PASCs, on average 18,470 per tissue (15%) had *wi*_1_*/wi*_2_ *>* 10, while DESeq2 analysis has identified a significant difference between read coverage in *wi*_1_ and *wi*_2_ for on average 43,615 (35%) of PASCs per tissue. In each tissue, on average 90% of PASCs with *wi*_1_*/wi*_2_ *>* 10 were also significant according to DESeq results. Since the results of the two methods overlapped, we chose to call a PASC with *wi*_1_*/wi*_2_ *>* 10 as expressed in the corresponding tissue.

We next compared the set of expressed PASCs to a reference set containing 689,346 PASs in 3’-UTRs of human genes that was derived from the GTEx using DaPars (*34*). Since the exact positions of PASCs in 3’-UTRs may vary, we selected 3’-UTRs that contain at least one expressed PASC and matched them against 3’-UTRs that were called as expressed by DaPars in genes with more than one annotated 3’-UTR. On average 85% of 3’-UTRs containing an expressed PASC were also called as expressed by DaPars, and vice versa 50% of 3’-UTRs called as expressed by DaPars contained at least one expressed PASC. That is, the expression of PASCs in tissues as measured by the *wi*_1_*/wi*_2_ metric and the results obtained by DaPars are consistent on a subset of PASCs in 3’-UTRs.

Previous studies have extensively characterized tissue-specific polyadenylation in the GTEx dataset using coverage-based methods, however implicitly focusing on polyadenylation in 3’-UTRs (*28, 34–36*). Here, we specifically considered intronic PASCs (iPASCs) identified by using polyA reads and examined the relationship between IPA and AS by juxtaposing the information on polyA reads to identify PASC positions, metrics based on split reads to measure AS rate, and the read coverage to assess IPA rate.

### Intronic polyadenylation and splicing

According to (*9*), alternative terminal exons that are generated through IPA can be categorized into two classes, skipped terminal exons (STE), which may be used as terminal exons or excluded, and composite terminal exons (CTE), which result from CPA in a retained intron (Figure 5A, right). To distinguish between these possibilities, we estimated the average read coverage in two additional windows, *we*_1_ and *we*_2_, at the exon-intron boundary (Figure 5A, left). For simplicity, the read coverage values in the four windows will also be denoted by *we*_1_, *we*_2_, *wi*_1_, and *wi*_2_. We expect that, in addition to large *wi*_1_*/wi*_2_ ratio, STE must be characterized by large *we*_1_*/we*_2_ ratio, while CTE must be characterized by small *we*_1_*/we*_2_ ratio.

**Figure 5:**
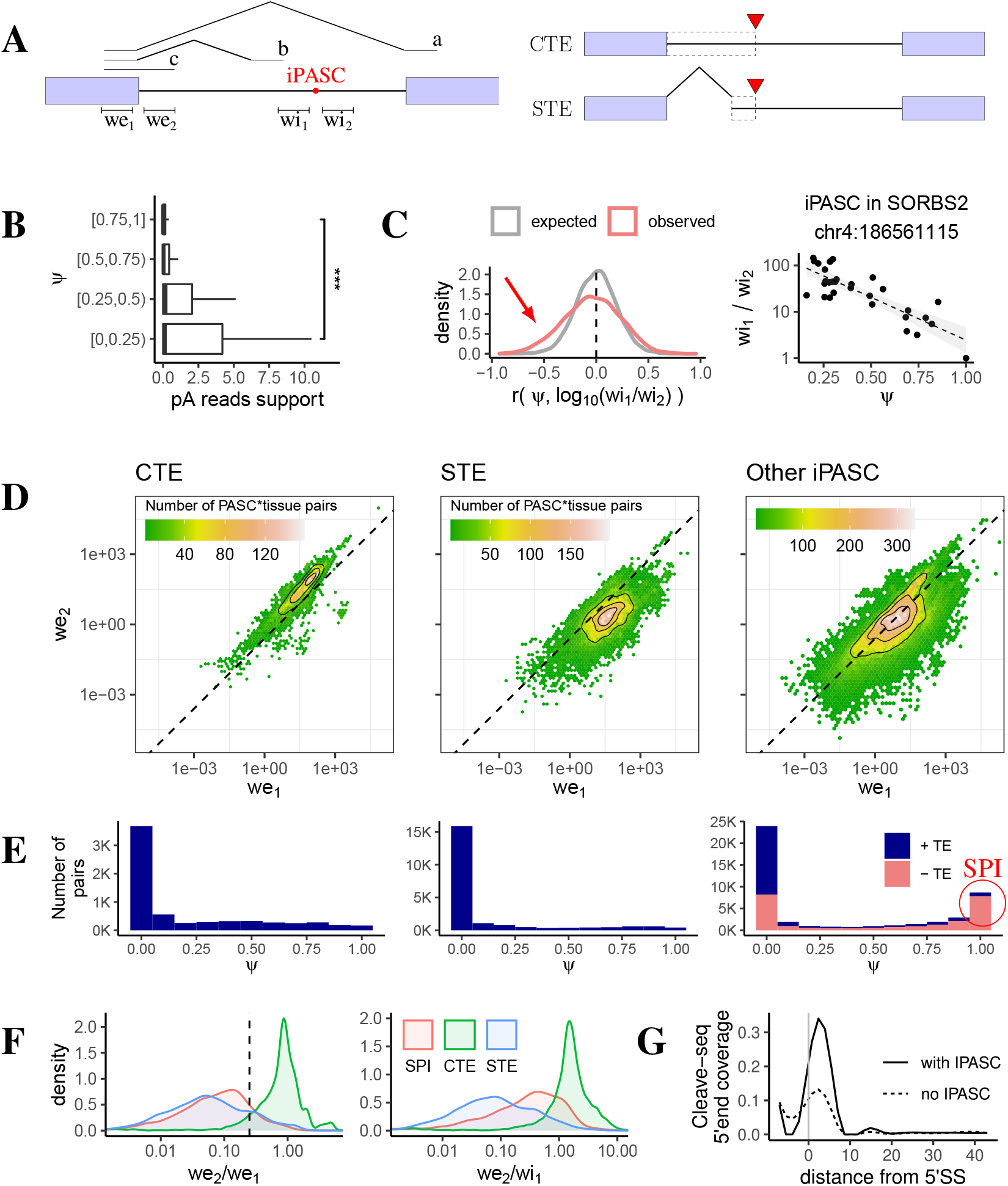
Intronic polyadenylation and splicing. **(A)** Exonic (*we*_1_ and *we*_2_) and intronic (*wi*_1_ and *wi*_2_) 150-nt windows. **(B)** The polyA read support of iPASCs in four *ψ* quartiles; *** denotes the 0.1% significance level. **(C)** Pearson correlation coefficients of *ψ* and log_10_(*wi*_1_*/wi*_2_) for *n* = 5, 081 iPASCs compared to the label-shuffled control (left). Negative association between *ψ* and log_10_(*wi*_1_*/wi*_2_) in the *SORBS2* gene. **(D)** Bivariate distribution of *we*_1_ vs. *we*_2_ in PASC-tissue pairs for CTE (*n* = 1, 342), STE (*n* = 2, 324), and other iPASCs (*n* = 59, 790). The dashed line corresponds to *we*_2_*/we*_1_ = 0.25. **(E)** The distribution of *ψ* for CTE, STE, and other iPASCs; +TE (*−* TE) denote iPASCs within (not within) 100 nts of an annotated TE. **(F)** The distribution of *we*_2_*/we*_1_ (left) and *we*_2_*/we*_1_ (right) values for CTE, STE, and SPI. The vertical dashed line denotes *we*_2_*/we*_1_ = 0.25. **(G)** The Cleave-seq 5’-end coverage in introns with (*n* = 33, 677) iPASC and without iPASC (*n* = 187, 531) under *XRN2* knockdown (see Figure S9D for the wild type).

To quantify the rate of splicing, we computed the number of split reads starting at the intron 5’-end and landing before iPASC (*b*), after iPASC at the canonical 3’-splice site (*a*), and the number of continuous reads (*c*) that span the exon-intron boundary (Figure 5A, left). These metrics were combined into the *ψ* = *a/*(*a* + *b* + *c*) ratio, referred to as the rate of canonical splicing, where *ψ* ≃ 1 indicates that the canonical splicing (*a*) prevails, while *ψ* ≃ 0 indicates the presence of AS events before iPASC. We expect that both STE and CTE are characterized by *ψ* ≃ 0 due to the lack of canonical splicing, with prevailing *b* in the case of STE and prevailing *c* in the case of CTE.

The values of *we*_1_, *we*_2_, *wi*_1_, *wi*_2_, and *ψ* were computed for 1,967,136 iPASC-tissue pairs comprising 63,456 iPASCs in 31 tissues. We observed a significant negative association between *ψ* and IPA rate measured by polyA read support (Figure 5B) or log_10_(*wi*_1_*/wi*_2_) (Figure S9A). This association also manifested itself as a negative skew in the distribution of Pearson correlation coefficients of *ψ* and IPA rate across tissues as compared to the background distribution, in which the tissue labels were shuffled (Figure 5C, left, and Figure S9B). The read coverage at iPASC changed two orders of magnitude when *ψ* increased from 25% to 100% in some remarkable cases (Figure 5C, right).

Further, we considered 88,856 iPASC-tissue pairs with a substantial read coverage drop at iPASC (*wi*_1_*/wi*_2_ *>* 10) and a substantially high read coverage in the upstream intronic window (*wi*_1_ *>* 0.1*we*_1_). The bivariate distributions of log(*we*_1_) and log(*we*_2_) for 1,342 annotated CTEs and 2,324 annotated STEs were separated by the line *we*_2_ = 0.25*we*_1_, with the former expectedly clustering above, and the latter clustering below the line (Figure 5D, left and middle). iPASCs other than CTE or STE formed a mixture of the two distributions (Figure 5D, right). A similar pattern was observed for the bivariate distributions of log(*wi*_1_) and log(*we*_2_) (Figure S9C). However, while *ψ* values of CTE and STE were characterized by a single peak at *ψ* ≃ 0 indicating the absence of canonical splicing (Figure 5E, left and middle), the *ψ* values of iPASC other than CTE or STE had a pronounced second peak at *ψ* ≃ 1 formed mostly by iPASCs without the TE support (Figure 5E, right). This peak is incompatible with CTE and STE models because it implies that IPA coexists with the canonical splicing. To further clarify this, we focused on iPASCs with *ψ >* 0.9, termed here as Spliced Polyadenylated Introns (SPI), and compared *we*_2_*/we*_1_ and *we*_2_*/wi*_1_ distributions among STE, CTE, and SPI (Figure 5F).

Similarly to STEs, SPIs were characterized by a low coverage in the intron 5’-end relative to the exon, yet a sufficiently high coverage upstream of iPASC relative to the intron 5’-end. We hypothesized that iPASCs with *ψ* ≃ 1 represent prematurely polyadenylated and spliced introns and hence expected them to have a monophosphate at the 5’-end (5’-p) resulting from the branchpoint (BP) cleavage by RNA debranching enzyme *Dbr1* (*45,46*). Then, the linearized product of CPA at an iPASC upstream of BP would consist of two separate molecules, one corresponding to the intronic RNA upstream of PAS with both 5’-p and polyA tail, and the other corresponding to the intron part downstream of PAS. Consistently with this, the 5’-end coverage of RNAs identified by Cleave-seq, a method designed to capture 3’-polyadenylated RNAs with 5’-p (*47*), was substantially larger in introns with iPASCs than in introns without iPASCs (Figure 5G) and, among the former, it was the largest in SPI (Figure S10). A similar enrichment in the 5’-end was also observed in 3’-pull down *in vitro* capping experiments (Figure S9E). Taken together, these results indicate that SPIs undergo both splicing and CPA and are not 3’-ends of distinct Pol II transcripts initiated and terminated within the same intron.

Next, we followed up a few cases of tissue-specific splicing and CPA (Figure 6). The iPASC in the *MEGF8* gene, which encodes a membrane protein associated with Carpenter syndrome (*48*), is an example of a CTE supported by intronic read coverage in absence of AS before PASC, most remarkably in thyroid tissue (Figure 6A). In the Attractin (*ATRN*) gene, which encodes a transmembrane protein associated with kidney and liver abnormalities in mice (*49*), an iPASC is expressed in muscle along with the elevation of read coverage in *wi*_1_ and activation of a splice site at its border, likely representing an unannotated STE (Figure 6B). Both these iPASCs are supported by *CSTF2* eCLIP peaks and evidence from PolyASite 2.0 (*24*). In contrast, iPASC in the *ATRX* gene, which encodes a chromatin remodeler linked to a range of diseases (*50*), exhibits elevated read coverage in *wi*_1_, but it lacks AS events that could support STE, or RNA-seq reads in the beginning of the intron that could support CTE (Figure 6C). Further manual inspection confirmed that the lack of split reads in this region is not a result of a mapping artifact. The only possible explanation for it would be that the canonical splicing and IPA co-exist and operate concurrently resulting in SPI.

**Figure 6:**
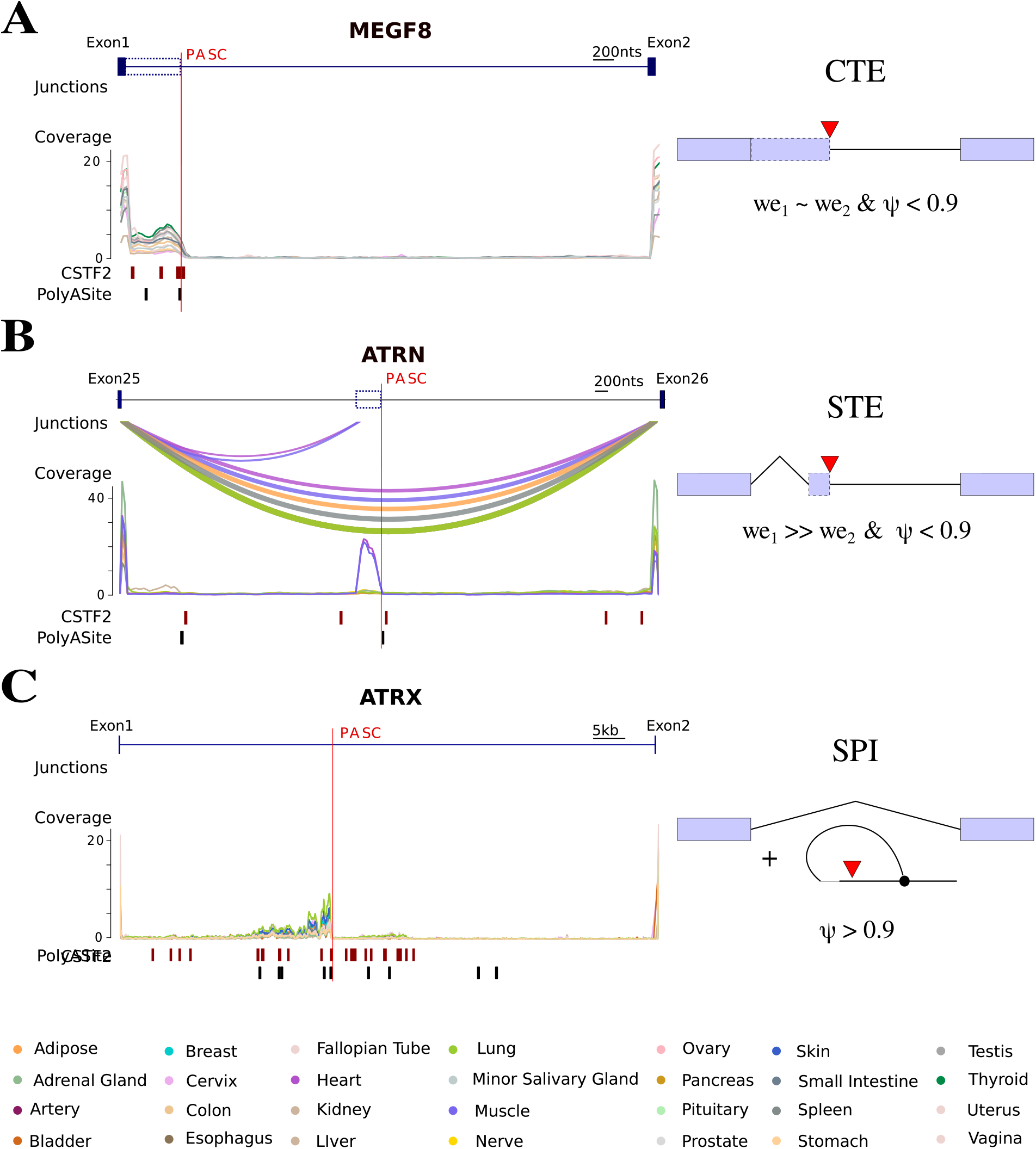
Case studies. **(A)** The iPASC between exons 1 and 2 of *MEGF8* generates a CTE. The eCLIP peaks of *CSTF2* and PASC from PolyAsite 2.0 are indicated in the track below. Arcs represent tissue-specific AS. **(B)** The iPASC between exons 25 and 26 *ATRN* generates a STE with tissue-specific expression in heart and muscle. **(C)** The iPASC between exons 1 and 2 likely generates a SPI because the intron 5’-end is not covered, *wi*_1_ is covered, but there is no evidence of STE.

To estimate the abundance of SPI events, we considered a strict set of iPASC-tissue pairs described above and categorized them as CTE, STE, and SPI according to the following criteria: *ψ* ≤ 0.9 and *we*_2_ *>* 0.25*we*_1_ (CTE), *ψ* ≤ 0.9 and *we*_2_ ≤ 0.25*we*_1_ (STE), and *ψ >* 0.9 (SPI) (Supplementary File 4). This yielded 3,352, 2,751 and 1,656 iPASCs corresponding to STE, CTE and SPI, respectively, where 58% of SPI were also supported by PolyASite 2.0 (*24*). We conclude that SPIs constitute a considerable fraction of IPA events, which are identified by polyA reads and 3’-end sequencing methods, and they must contribute greatly to the observed landscape of intronic polyadenylation.

## Discussion

Thousands of recurrent and dynamically changing IPA events have been identified by 3’-end sequencing methods, but the matched data to study the interplay between IPA and AS in the same biological condition are currently in high demand (*11*). The GTEx dataset represents an ideal resource for studying this interplay because the information on the positions and tissue-specific expression of intronic PASs, which is captured by polyA reads, is complemented by tissue-specific splicing rates inferred from split reads that align to splice junctions. In this work, we used for the first time the approach based on polyA reads, one that was applied previously to much smaller datasets (*30, 31*), for the identification of PASs at the scale of GTEx project and combined it with a coverage-based method to assess tissue-specific IPA rates.

The admixture of polyA reads in a large array of RNA-seq experiments provided an invaluable source of information about CPA. However, this approach may have limitations related to the mappability of reads with long soft clip regions as the frequency distribution of polyA reads decays with increasing the length of the overhang. The positional distribution of PASCs in constitutive exons and introns has a pronounced peak in the end of exonic and in the beginning of intronic regions (Figure S5) resembling clusters of CAGE tags near internal exons and occurrence of polyA-seq peaks close to exon boundaries (*51, 52*). These anomalies likely arise from erroneous mappings of split reads that contain the polyA tail, e.g. when the adenine-rich part of the read or a short segment between splice junction and the stretch of non-templated adenines are incorrectly attributed to soft clip region (example in Figure S7). However, these details do not invalidate the polyA-based method since the positional distribution of PASCs obtained by other protocols, e.g., in PolyASite 2.0, has similar peaks near exon boundaries (Figure S6) indicating that alignment of split reads with short exonic parts is a common problem of all such methods.

The widespread nature of IPA has been appreciated recently with the development of 3’-end sequencing (*53*). Functionally important IPA cases have been described in specific genes (*10, 54–58*), however most transcripts harboring incomplete reading frames translate into potentially deleterious, truncated proteins that may pose a hazard to the cell (*59*). In eukaryotes, they usually lack normal termination codons and are rapidly degraded via nonsense mediated decay or nonstop decay pathways (*60, 61*). We found that the majority of polyA reads align to 3’-UTRs, but a sizable fraction (5–8%) still map to the coding part. Intriguingly, PASs within the coding part appear to be more frequent in introns than in exons, which is partly explained by the higher GC content and stronger evolutionary constraints against generating the canonical AATAAA consensus sequence in exons. However, a remarkably large number of intronic PASs raises concerns about their implication in premature transcription termination (*62*) and hints at the existence of a mechanism that counteracts their activity. How could it be that 87% of human protein-coding transcripts contain an intronic PAS, but cells are still able to produce full-length transcripts?

Here, we speculate that a sizable fraction of intronic PASs observed in polyA read analysis (and also in 3’-end sequencing) represent SPIs, which are generated by splicing and CPA operating concurrently with each other and with the elongating transcription (Figure 7). The spliceosome and the CPA machinery operate at their intrinsic rates that are subordinate to the transcription elongation speed, and one of them could be faster than the other depending on the biological condition. If CPA occurs faster, then it will lead to the generation of a truncated transcript with CTE. If splicing happens faster, then the intron containing PAS will be spliced out, and PAS will be degraded as a part of the lariat. However, if the spliceosome has already assembled on the intron when CPA started PAS-mediated cleavage, the second catalytic step of the splicing reaction will remove the cleaved and polyadenylated part, resulting in SPI (Figure 7 middle). Consequently, SPIs are intronic RNAs spanning from the 5’-splice site to PAS that contain both 5’-p resulting from lariat debranching by *Dbr1* and the polyA tail. Being transient byproducts of the coupling between splicing and CPA, they must be degraded from the 5’-end by cellular exonucleases, which is evidenced in many cases as a characteristic noisy ramp in the read coverage that gradually increases from the 5’-splice site to PAS (Figure 6C). Nonetheless, the intermediates of SPI digestion are still visible as polyA reads due to the presence of the polyA tail. Our conservative estimates indicate that SPIs constitute almost a quarter of all IPA events, and more than half of SPIs are supported by PAS from PolyASite 2.0 (*24*). We conclude that 3’-end sequencing methods may overestimate the rate of IPA, and that their results require careful interpretation.

**Figure 7:**
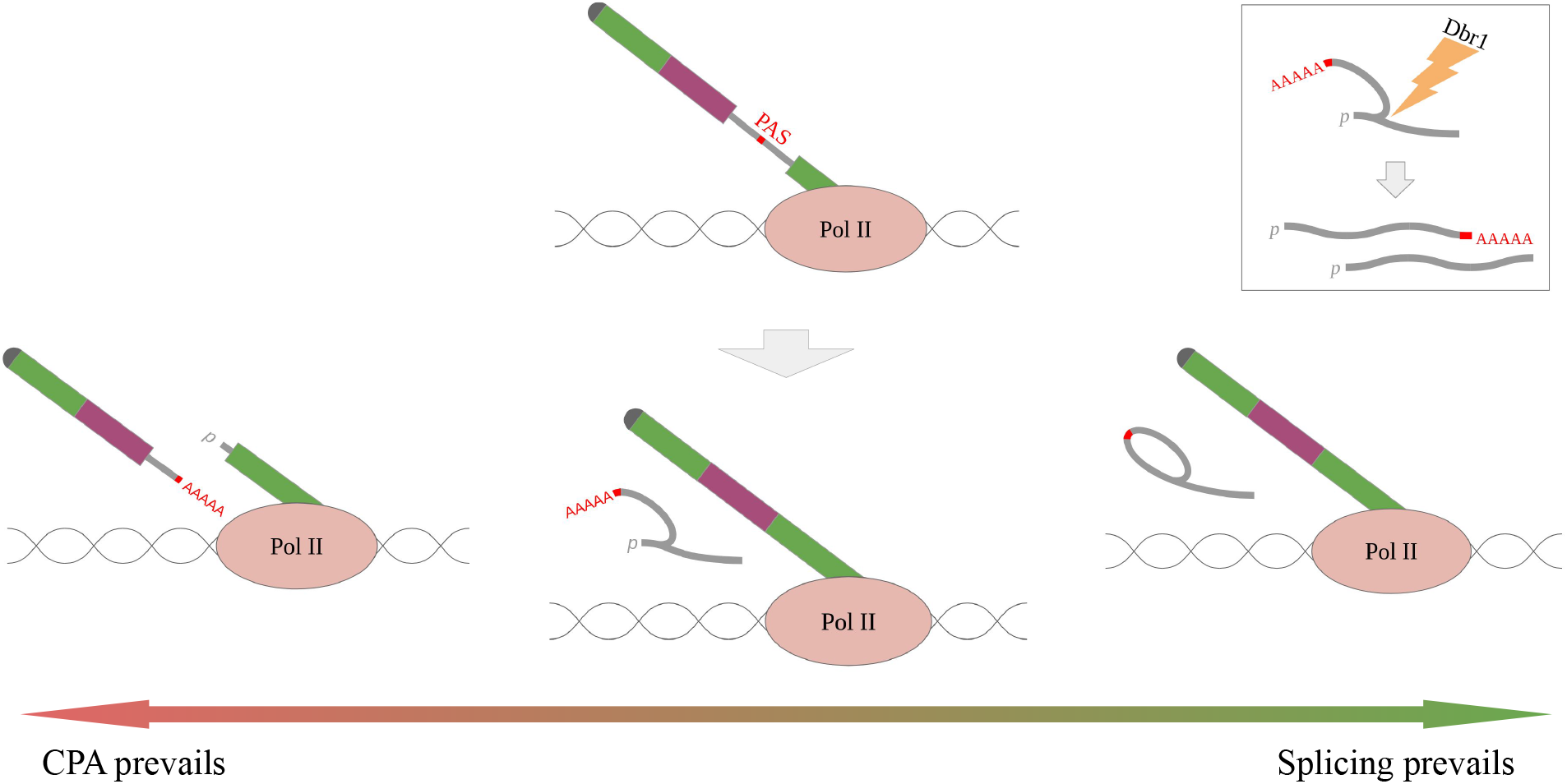
Spliced Polyadenylated Intron (SPI). When the CPA rate exceeds the splicing rate, IPA leads to the generation of a truncated transcript isoform (left). When the splicing rate exceeds the CPA rate, the intron is spliced out and PAS is degraded as a part of a lariat (right). When the CPA and splicing machinery operate at the same rate, the intron is cleaved and polyadenylated while it is being spliced (middle) resulting in SPI. The lariat is debranched producing two separate RNAs (inset) corresponding to intron fragments upstream and downstream of PAS, where the upstream part contains both 5’-p and polyA tail.

In this light, it appears plausible that a side function of co-transcriptional splicing could be to rescue transcripts from premature transcription termination by dynamically counteracting CPA. This hypothesis challenges the current assumption that when an intronic PAS is used, the intron is not spliced and *vice versa*. Temporal and spatial interactions of splicing and CPA are orchestrated by a multitude of factors playing dual roles, which recognize signals that are located in the nascent pre-mRNA and bind the same pre-mRNA substrate at the same time (*14, 63, 64*). It is therefore not impossible that evolution allowed for the generation of dispensable intronic PAS, which are spliced out co-transcriptionally and manifest themselves in the 3’-end sequencing data as SPIs. Whether or not SPIs are functional on their own is a matter of further investigation.

## Conclusion

Massive amounts of RNA-seq data in the GTEx dataset offered a unique possibility to analyze tissue-specific splicing and polyadenylation. The observed patterns of intronic polyadenylation and splicing reconfirm that splicing and polyadenylation are two inseparable parts of one consolidated pre-mRNA processing machinery, leading to the conjecture that co-transcriptional splicing is a natural mechanism of suppression of premature transcription termination.

## Methods

### Genome assembly and transcript annotation

February 2009 (hg19) assembly of the human genome and GENCODE transcript annotation v34lift37 were downloaded from Genome Reference Consortium (*65*) and GENCODE website (*43*), respectively. Transcript annotations were parsed by custom scripts to extract the coordinates of transcript ends, exons and introns. The attribution of PAS to protein-coding, non-coding, and intergenic segments was done on the basis of their occurrence in the corresponding gene types.

### Partition of protein-coding genes

To partition protein-coding genes into segments, we parsed the annotation of protein-coding transcripts from GENCODE and extracted 5’-UTRs, 3’-UTRs and CDS of all transcripts as follows. Genomic regions that were not covered by any transcript were classified as intergenic. A gene part was classified as 5’-UTR (respectively, 3’-UTR) if it belonged to the 5’-UTR (respectively, 3’-UTR) of at least one annotated transcript of the gene; the rest of the gene sequence was classified as CDS. We next considered exons and introns of all annotated protein-coding transcripts and used them to further subdivide CDS regions into exonic, intronic, and alternative regions. A genomic region was classified as always exonic (respectively, intronic) if it belonged to exonic (respectively, intronic) parts of all annotated transcripts that overlap the region; otherwise, it was classified as an alternative exonic region.

### Identification of PAS from RNA-seq data

GTEx RNA-seq data were downloaded from dbGaP (dbGaP project 15872) in fastq format and aligned to the human genome assembly hg19 using STAR aligner version 2.7.3a in paired-end mode (*66*). PySAM suite was used to extract uniquely mapped reads (NH:1) (*67*). To identify polyA reads, we considered all reads containing a soft clipped region of at least 6 nts excluding reads with average sequencing quality below 13, which corresponds to the probability 0.05 of calling a wrong base. We required that the reported nucleotide sequence of the clipped region, which always corresponds to the positive strand according to BAM format, contained at least 80% T’s if the soft clip was in the beginning of the read, and 80% A’s if the soft clip was in the end of the read. In fact, the requirement of 80% A’s or T’s was excessively strict since 87% of soft clip regions consisted entirely of A’s or T’s. Samples that contained an exceptionally high number of polyA reads were excluded from analysis (Figure S1). PolyA reads were pooled by the genomic position of the first non-templated nucleotide, referred to as PAS position, resulting in read counts (*f*_*i*_) for each value of the overhang (*i*). Accordingly, each PAS was characterized by the number of aligned polyA reads

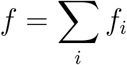

and Shannon entropy of the overhang distribution

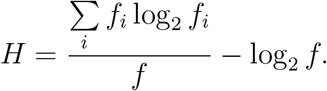

In order to select a reasonable number of PAS, we repeated the above steps using an array of thresholds on the minimal overhang length and Shannon entropy threshold *H* and computed the number of annotated gene ends that are supported by PAS (Figure S2). The threshold *H* ≥ 2 in combination with the minimum overhang length of 6 nts appears to be optimal since it captures 85% annotated gene ends and yields 565,387 PAS, a number that corresponds by the order of magnitude with the size of the PAS set reported in PolyASite 2.0 (*24*). PASs that were located within 10 nts of each other were merged into clusters (PASCs) using the GenomicRanges package (*68*).

### Precision and recall

The list of PASCs obtained from the GTEx RNA-seq data (referred to as GTEx) was validated against two reference sets, the published set of PASCs inferred from the 3’-end sequencing (*24*) (referred to as Atlas) and the set of annotated TEs provided by GENCODE consortium (*43*) (referred to as GENCODE). In each comparison, we calculated the precision and recall metrics of GTEx with respect to the reference set by imposing variable thresholds on PASC support level. First, GTEx and Atlas were both compared to GENCODE so that a PASC was considered a true positive if it was located within 100 nts from an annotated TE. The precision and recall metrics varied depending on the number of supporting polyA reads (in GTEx) and the average expression (in Atlas) reaching the optimal *F*_1_ = 2(*P −*^1^ + *R*^*−* 1^) −^1^ score at *P* = 0.57 *−* 0.58 and *R* = 0.49− *−* 0.51 (Figure S3, top left). The same scores, in which each PASC was weighted by the read support, showed a similar performance with the optimal *F*_1_ score of *P* = 0.83 *−* 0.86 and *R* = 0.73 − 0.76 (Figure S3, bottom left). In comparison to Atlas as a reference set by the number of PASC, GTEx showed a moderate performance with *P* = 0.66 and *R* = 0.30, especially in terms of recall, i.e., a large fraction of PASCs from Atlas were not detected (Figure S3, top right). However, when the same comparison was made by the number of transcripts, i.e., by weighting PASCs by the read support, the precision and recall were 0.92 and 0.97, respectively, indicating that the GTEx primarily misses PASCs with low level of read support (Figure S3, bottom right).

### Relative position in the gene

For each PASC, which is characterized by the interval [*x, y*] in the gene [*a, b*], where *x, y, a*, and *b* are genomic coordinates on the plus strand, we defined *p*, the relative position in the gene as 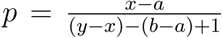 for genes on the positive strand, and used the value of 1 *− p* for genes on the opposite strand. The values of *p* outside of the interval [0, 1] indicate that the PASC is located outside of the annotated gene boundaries. In the same way, PASC relative positions were computed in exonic and intronic regions.

### Read coverage and fold change

To quantify the extent, to which CPA happen at a specific PASC in a specific tissue, we first calculated the read coverage genomewide for each GTEx sample by considering only uniquely mapped reads (MAPQ=255 when processed via STAR mapper) with `bamCoverage` utility using flags -binSize 10 –minMappingQuality 255 (*69*) and averaged the read coverage values between samples within each tissue using `wiggletools mean` utility (*70*).

Next, we calculated the mean read coverage per nucleotide in 150-nt windows starting 10 nts upstream and downstream of each PASC in each tissue (referred to as *wi*_1_ and *wi*_2_) using `multiBigwigSummary utility` (*69*) and computed the log-fold-change metric as the logarithm of the ratio of the mean read coverage in the upstream and downstream windows, respectively, with a pseudocount of 10^−3^. To take into account the variation between samples when assessing PASC expression, we followed the approach described previously (*11*) by detecting significant differences in read counts between the upstream and downstream windows (*p*_*adj*_ *<* 10^−3^) using DESeq2 (*44*), separately in each tissue.

Intronic PASCs were defined as PASCs located within at least one annotated intron of a protein-coding gene *>*200bp away from the closest annotated splice site (*n* = 63, 456). The shortest intron containing a PASC was chosen, and the average read coverage was computed not only in *wi*_1_ and *wi*_2_, but also in 150-nt windows starting 10 nts upstream and downstream of the intron 5’-end (*we*_1_ and *we*_2_, Figure 5A). An intronic PASC located within 100 nts from an annotated TE of a protein-coding transcript (*n* = 3, 781) was categorized as STE (respectively, CTE) if the terminal exon of the transcript fully belonged to the containing intron (respectively, overlapped the interval from 5’-splice site to PASC). This categorization yielded 1,342 CTEs and 2,324 STEs, while 115 PASCs were located near TEs of several transcripts resulting in conflicting annotation.

To estimate the mean read coverage in constitutive exons, alternative exons, and introns, the total read coverage values per nucleotide in all GTEx samples were averaged between windows located in the respective regions to obtain normalization factors (3.3 · 10^6^, 3.2 · 10^6^, and 8.0 · 10^4^, respectively). The latter were used to normalize the fraction of polyA reads in the respective regions (Figure 3C) relative to the average read coverage.

### Splicing metrics

To quantify tissue-specific alternative splicing associated with intronic PASCs, we computed split read counts using the IPSA pipeline as explained earlier (*33, 71*). The counts of split reads were pooled within each tissue to compute the *ψ* = *a/*(*a* +*b* +*c*) metric (Figure 5A). The values of *ψ* with the denominator below 30 were discarded as unreliable.

### Cleave-seq 5’-end coverage and 3’-RNA capping and pulldown data

Cleave-seq data in HeLa cells were downloaded from Gene Expression Omnibus under the accession number GSE165742 (samples GSM5566266–GSM5566269) in bigwig format (*47*). The per-bin cleave-seq signal was computed around 5’-splice sites using deeptools computeMatrix tool with the following parameters `computeMatrix reference-point -a 150 -b 20 -bs 5 --nanAfterEnd --missingDataAsZero --skipZeros` and consequently averaged between replicas and introns for visualization.

The 3’-RNA capping and pulldown (3’-PD) data in U2OS cells (*72, 73*) were downloaded from Gene Expression Omnibus under the accession number GSE84068 including three 3’-PD replicas (GSM2226722–GSM2226724) and three total polyA^+^ RNA-seq replicas for normalization (GSM2226713–GSM2226715). The per-bin coverage around 5’-splice sites was computed as for Cleave-seq. For visualization, the 3’-PD coverage values were averaged between replicates, normalized to the respective total RNA-seq coverage in each bin of each intron, and averaged between introns.

## Availability of data and materials

The datasets generated during the current study are available online at https://zenodo.org/record/6587186. The source code used for the analysis is available at https://github.com/mashlosenok/RNAseq_PAS_finder.

## Authors’ contributions

DP designed and supervised the study; MV and SM performed data analysis; DP and MV wrote the first draft of the manuscript. All authors edited the final version of the manuscript.

## Acknowledgments

The authors thank Vera Rybko, Dmitry Skvortsov, and Olga Donstova for insightful discussions on molecular mechanisms of splicing and polyadenylation.

