## Supplementary Information for "Transcriptome sequencing suggests that pre-mRNA splicing counteracts widespread intronic cleavage and polyadenylation"

January 17, 2023

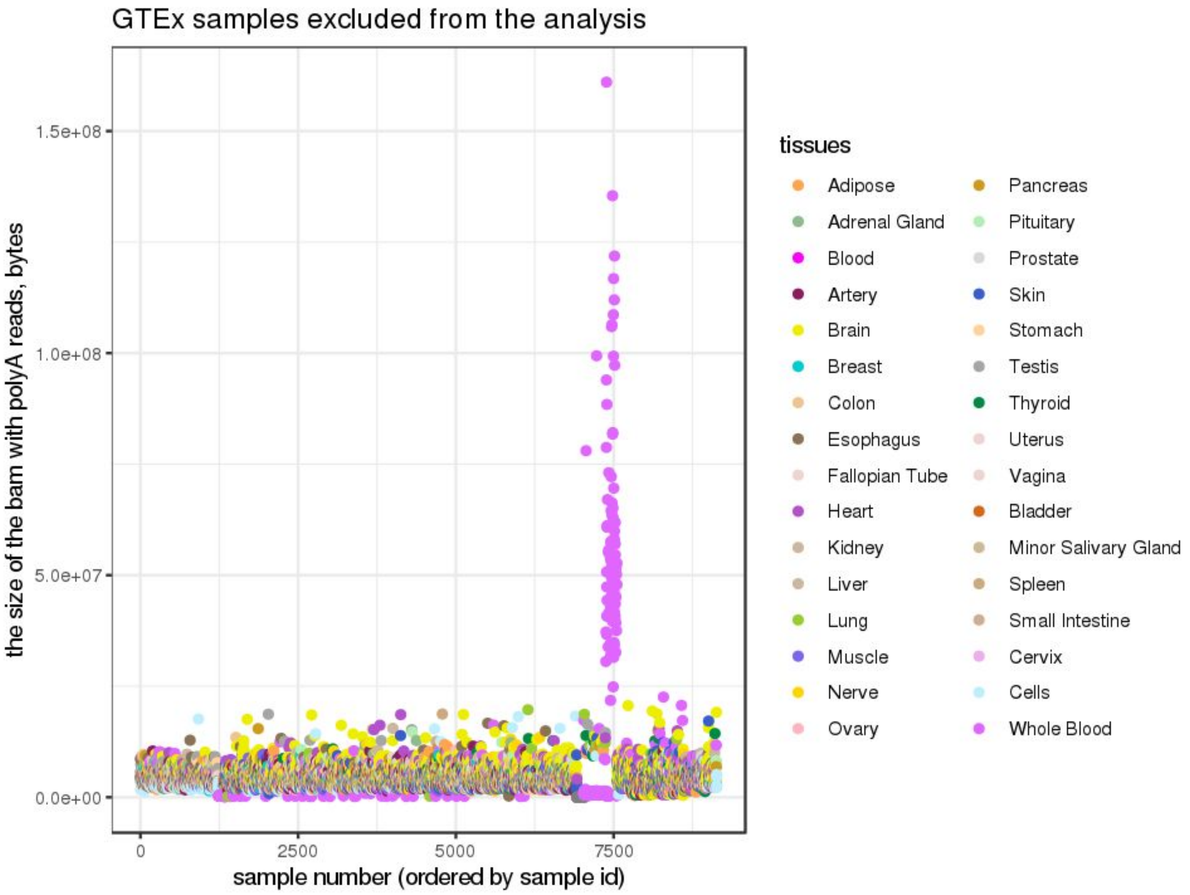

**Figure S1:** A diagram of the number of polyA reads (measured as BAM file size) of 9,135 GTEx samples, of which 220 samples had exceptionally large number of polyA reads and were excluded from further analysis. The excluded samples are from Whole Blood tissue type.

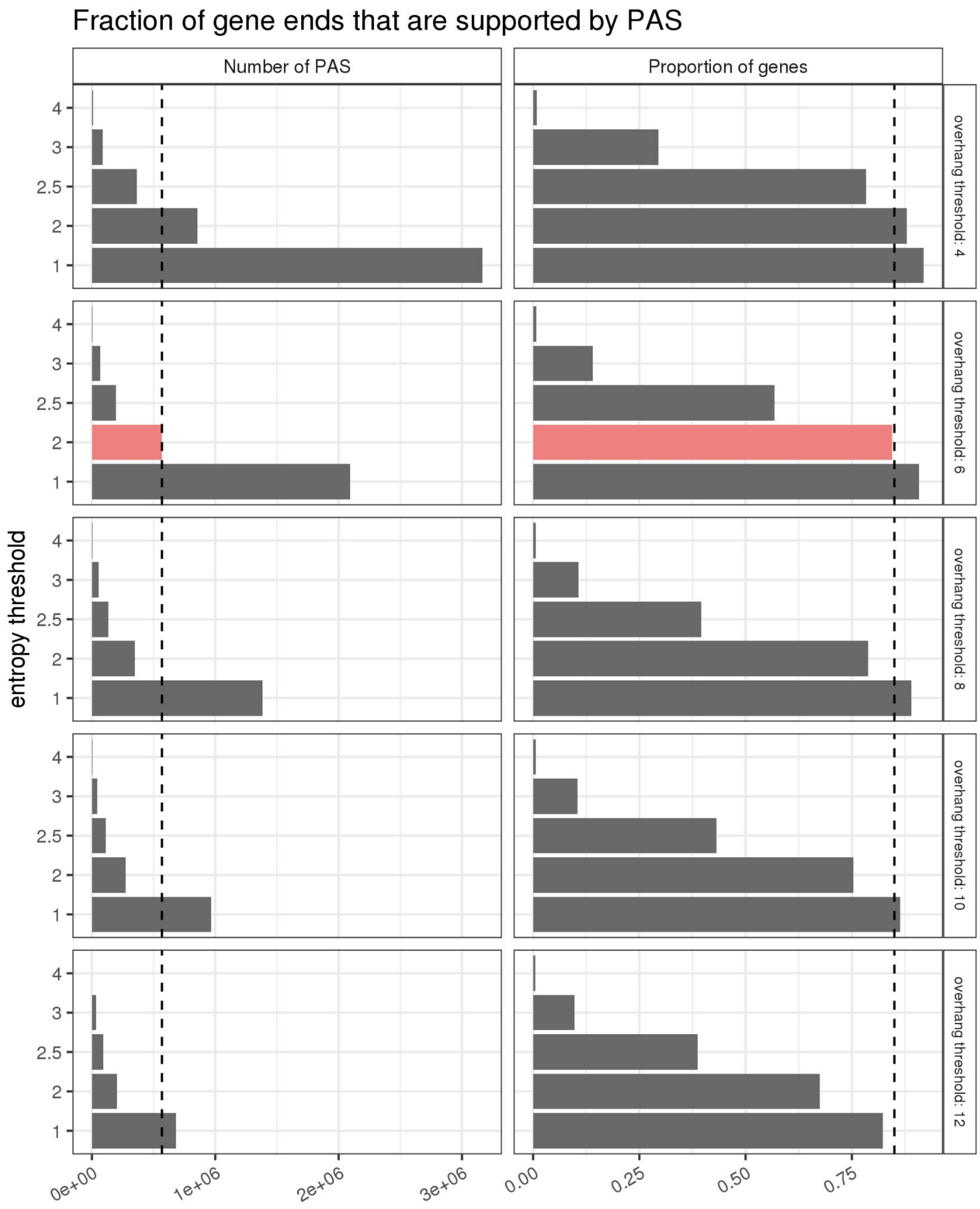

**Figure S2:** The PAS identification procedure was carried out using an array of thresholds on the minimal overhang length and Shannon entropy ($H$). Shown are the number of PAS (left) and the proportion of genes, in which an annotated transcript end was identified within 100 bp of a PAS. The dashed line represents the number of PAS reported in PolyASite 2.0. The condition $H \geq 2$ in combination with the minimum overhang length of 6 nts (red) gives the optimal cutoff.

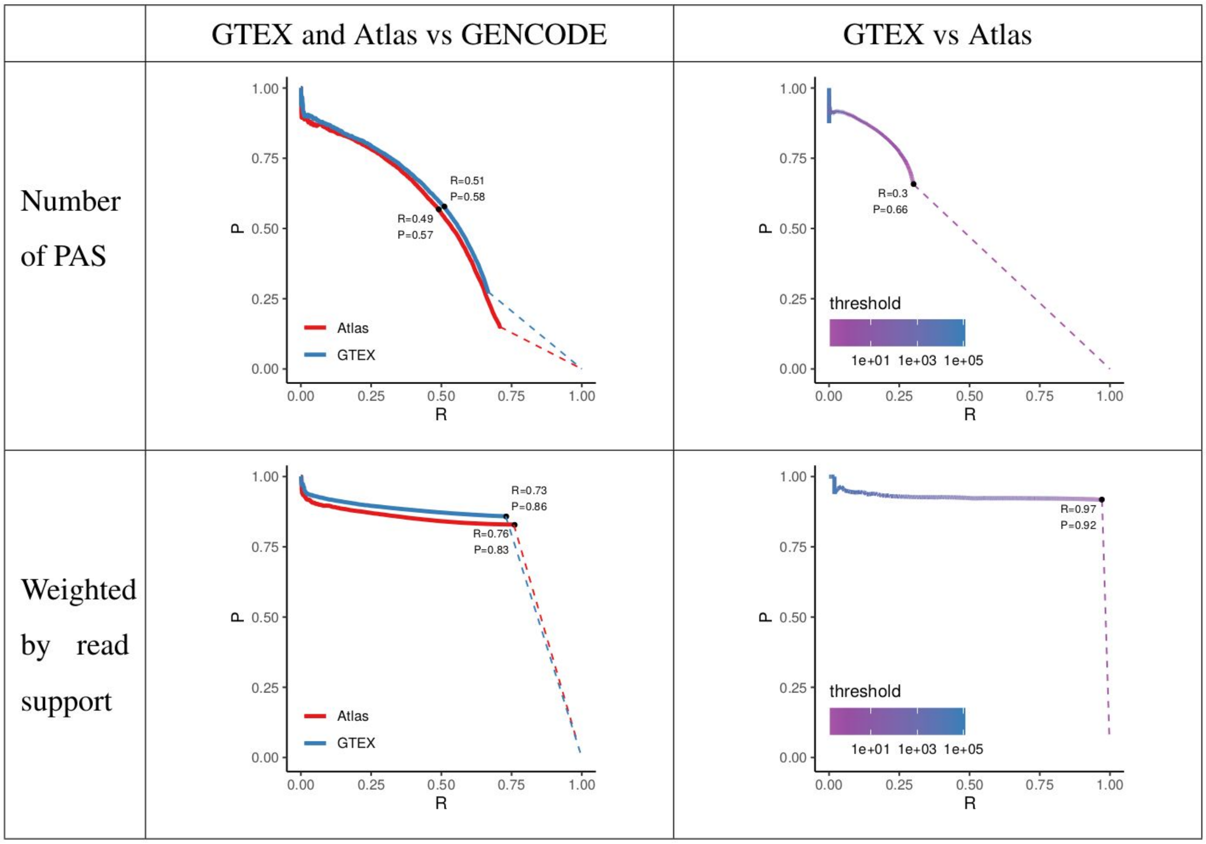

**Figure S3:** The validation of PASCs from GTEX and Atlas vs. GENCODE (left) and GTEX vs. Atlas (right) in terms of the number of PASCs (top) and the number of polyA reads (bottom) using precision ($P$) and recall ($R$) metrics (see Methods for details). The curves are obtained by varying the threshold on polyA read support. The values of precision and recall with the largest $F_1$ metric are shown.

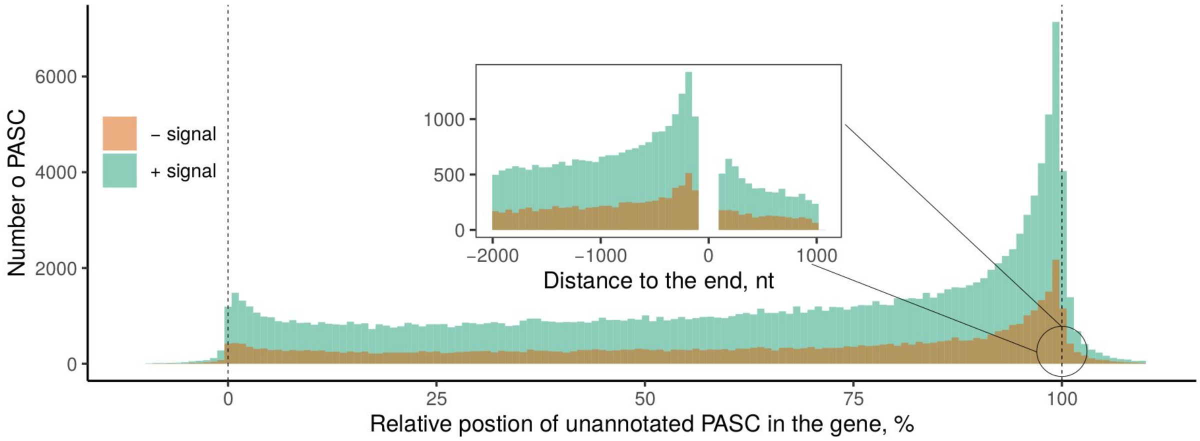

**Figure S4:** The relative positions of unannotated PASCs (i.e., ones not within 100 bp of any annotated TE) along the gene length for PolyASite 2.0. The inset shows distribution of absolute positions of unannotated PASCs around the gene end (see Figure 2E legend).

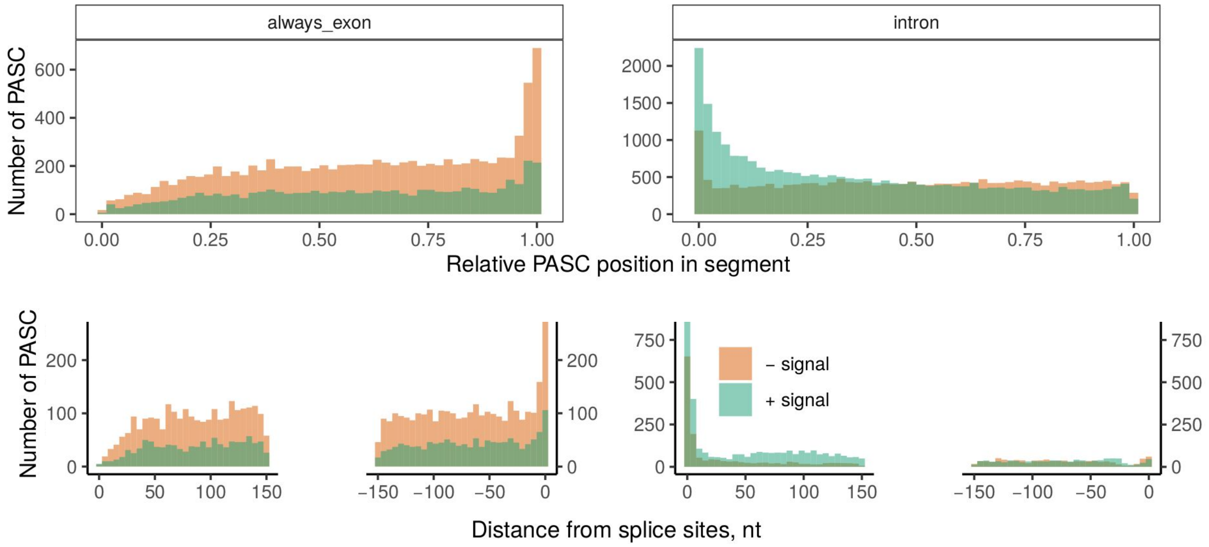

**Figure S5:** The relative (top) and absolute (bottom) positions of PASCs from GTEx in constitutive exons (always exon) and introns. The enrichment of PASCs at the end of constitutive exons and in the beginning of introns in part can be attributed to mapping artifacts (see Discussion) for PASCs with (+signal) and without a signal (−signal).

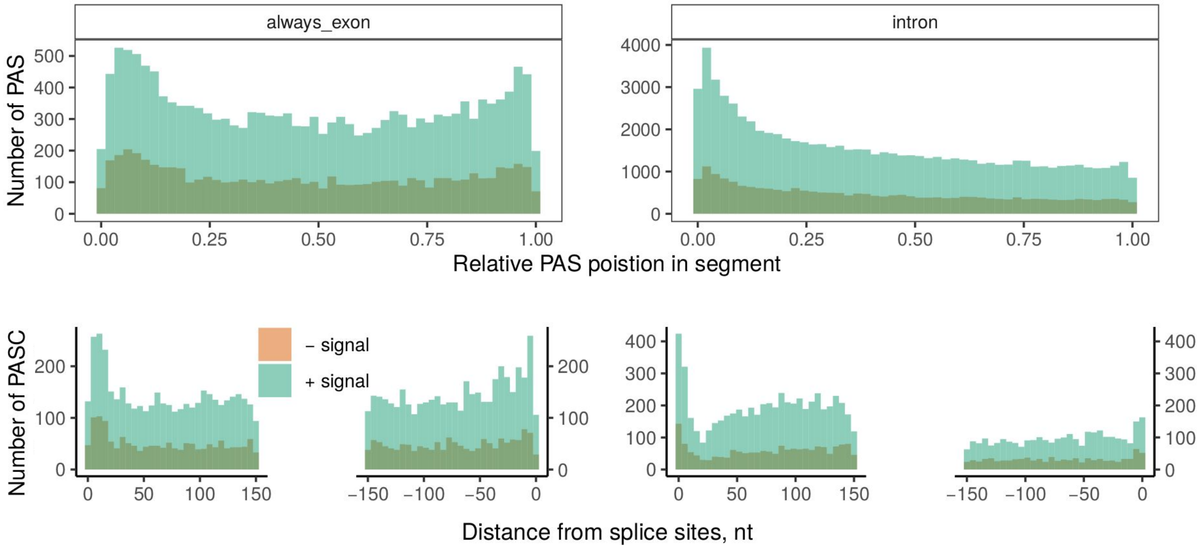

**Figure S6:** The relative (top) and absolute (bottom) positions of PASCs from PolyASite 2.0 in constitutive exons (always exon) and introns (see Figure S5 legend).

## A

polyA reads:

SRR1400910.12641453 83 9 19120877 255 8S68M 9 19119788 68
TTCTTTTTCTAGTAATTCCTCAATGAGAGGGAGGTACTGTTTTACCAACAGCAATGATTTGGTGAGTGCATTTACT

SRR1400910.12645416 163 9 19120877 255 6S70M 9 19120948 70
CTTTTTCTAGTCCTTCCTCAGTGAGAGGGAGGTACTGTTCTACCAACAGCTCTGATTTGGTGAGTGCATTTTCTAC

SRR1400910.12652323 163 9 19120877 255 9S67M 9 19123575 67
TTTTTTTTTCTAGTTCGTCCTCAGTGAGAGGGAGGTACTGTTCTACCAACAGCTCTGATTTGGTGAGTGCATTTTC

Corresponding region:

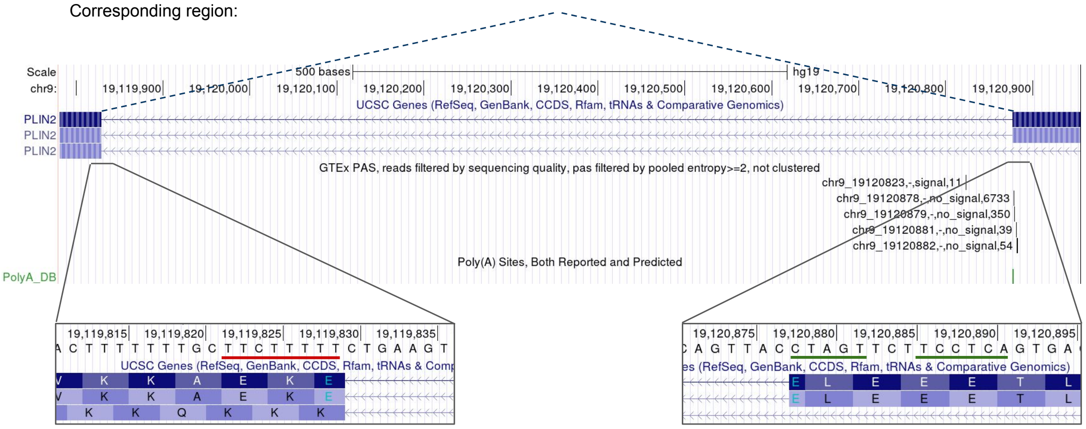

## B

polyA reads:

SRR1402414.19165097 163 14 23851636 255 8S68M 14 23851740 68
ATTTTTTTCTTGGCACCAATGTCACGGCTCTTGGCTCGAAGCTTGTTGACCTGGGACTCATCGATGTCCGCCCGCT

SRR1402414.19165230 147 14 23851636 255 7S69M 14 23851140 69
TTTTTTGCTTGGCCCCAATGTCACGGCTCTTGGCTCGAAGCTTGTTGACCTGGGACTCAGCGATGTCCGCCCGCTC

SRR1402414.19180339 163 14 23851636 255 6S69M1S 14 23851721 69
TTTTTGCTTGTTCACCAATGTCACGGCTCTTGGCTCGAAGCTTGTTGACCTGGGACTCAGCGATGTCCGCCCGCTCN

Corresponding region:

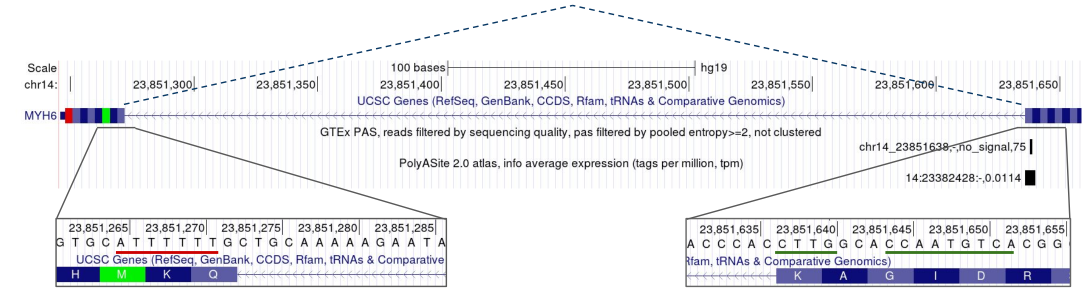

**Figure S7:** Two examples (A and B) of mapping artifacts, in which a spurious PAS is incorrectly placed near exon boundaries due to the presence of mismatches (underlined) followed by A-reach tracks. Short read alignments are shown on the positive strand as T-reach track (red) as they appear in BAM file. The soft clip region is shown in green.

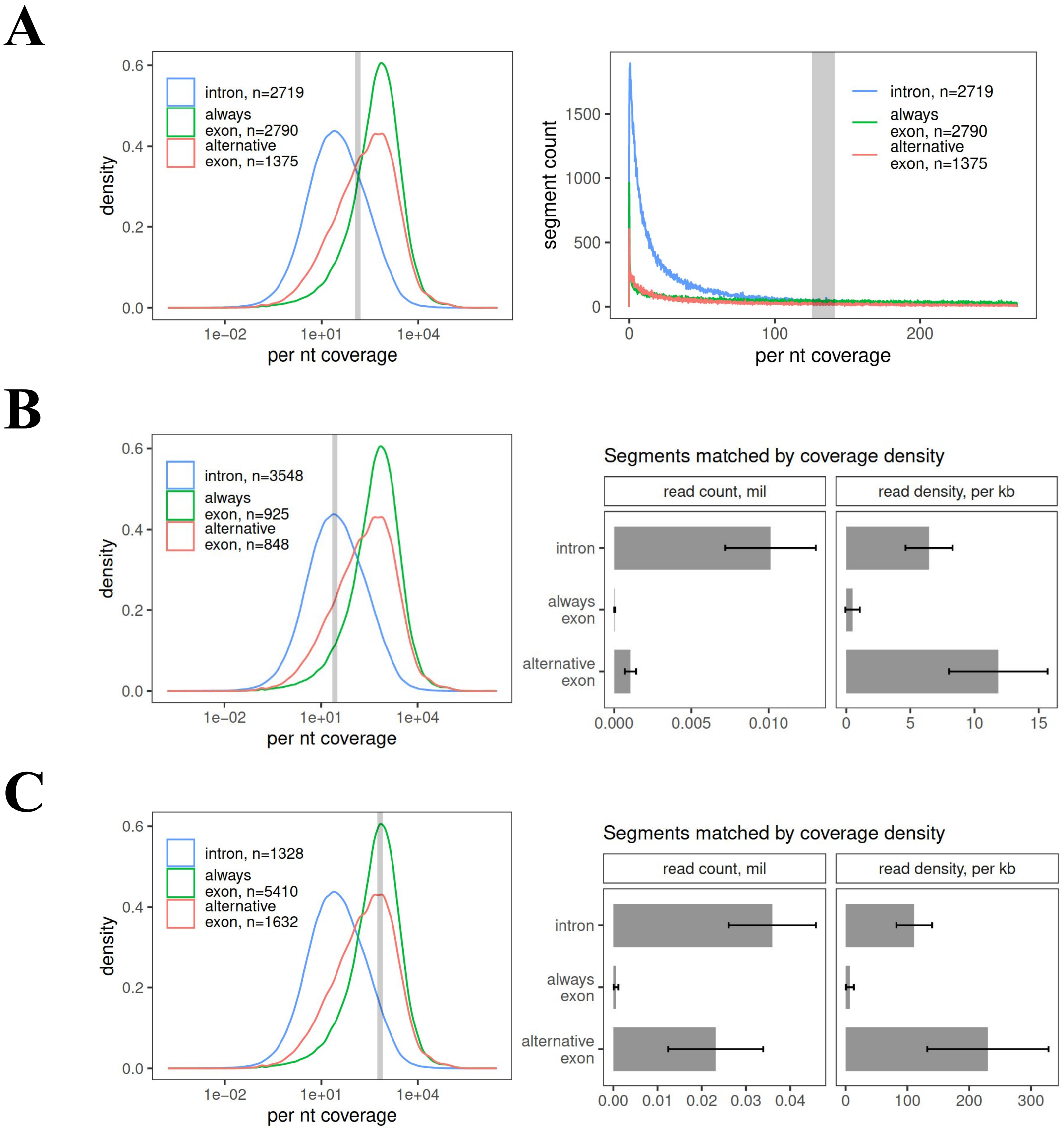

**Figure S8:** Matching genomic intervals by read coverage. (**A**) The distribution of read coverage density values in three types of genomic segments. For each segment, the mean read coverage was computed as the number of reads per kb per sample. The segments with average read coverage of $133 \pm 6.7$, which corresponds to the intersection of density curves (highlighted by grey area), were selected for the analysis in Figure 3D (approximately 1375 segments of each type). (**B**) The same procedure repeated for the average read coverage interval $27 \pm 5$ (left) and the distribution of polyA reads as in Figure 3D (right). (**C**) The same procedure repeated for the average read coverage interval $666 \pm 5$ (left) and the distribution of polyA reads as in Figure 3D (right).

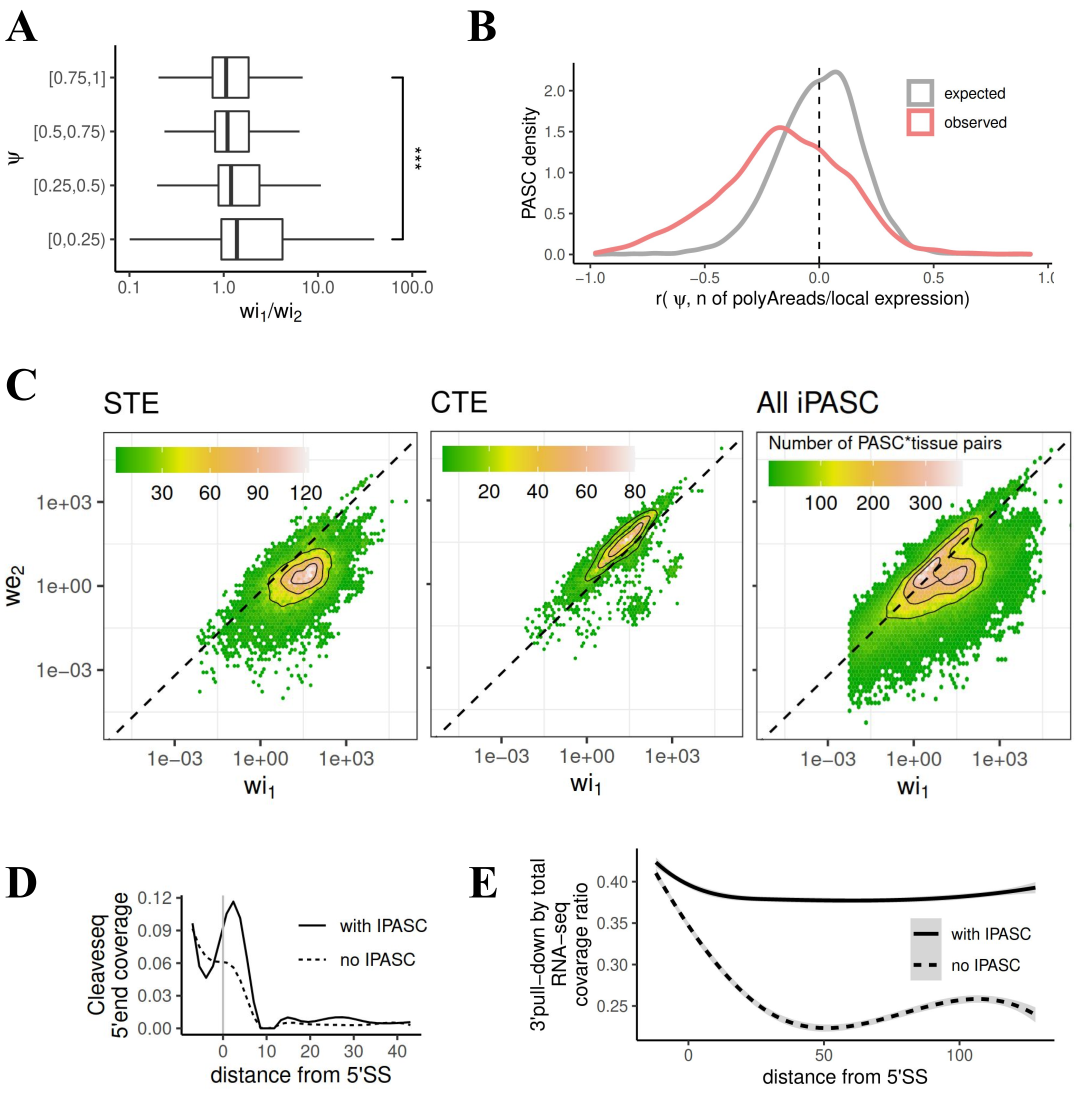

**Figure S9:** iPASC properties. (**A**) The distribution of $wi_1/wi_2$ values in four $\psi$ quartiles. The $wi_1/wi_2$ significantly increases with decreasing $\psi$. (**B**) The distribution of Pearson correlation coefficients between number of polyA reads normalized by number of reads spanning the 5' splice site and $\psi$. (**C**) The joint densities of $wi_1$ and $we_2$ for iPASCs that correspond to the annotated STE, CTE and for all expressed iPASC. (**D**) Cleaveseq 5'-end densities in introns with and without iPASCs (without *XRN2* knockdown). (**E**) Normalized coverage in 3'-pull down *in vitro* capping experiments as a function of the distance from the 5'-splice site, in introns with and without iPASCs.

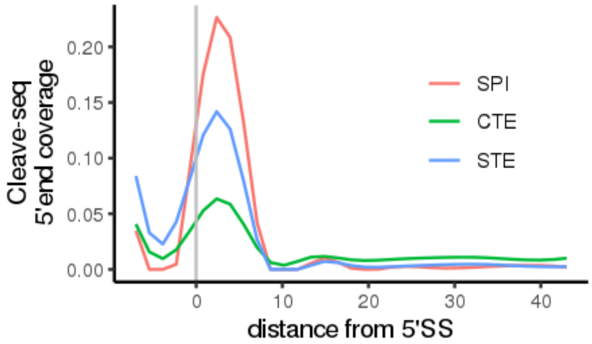

**Figure S10:** The Cleave-seq 5'-end coverage in introns with STE, CTE, and SPI under *XRN2* knockdown. Introns were categorized as containing CTE ($n = 247$), STE ($n = 360$), or SPI ($n = 262$) if they contained an unambiguous iPASC of the respective type, while introns with iPASCs of different types were discarded.

**SupplementaryDataFile 1:** The list of 565,387 PAS with $H \geq 2$ and minimum overhang of six nucleotides.

**SupplementaryDataFile 2:** The list of 318,898 PAS clusters in protein-coding genes.

**SupplementaryDataFile 3:** The list of 126,310 PASCs located >200 nts away from exon boundaries and their expression levels in GTEx tissues.

**SupplementaryDataFile 4:** The list of 63,456 intronic PASCs in 31 tissues, their expression levels in GTEx tissues, annotation status and categorization as CTE, STE, and SPI.
